## Supplemental Figure 1 for "Peculiar transcriptional reprogramming with functional impairment of dendritic cells upon exposure to transformed HTLV-1-infected cells"

### Slide 1
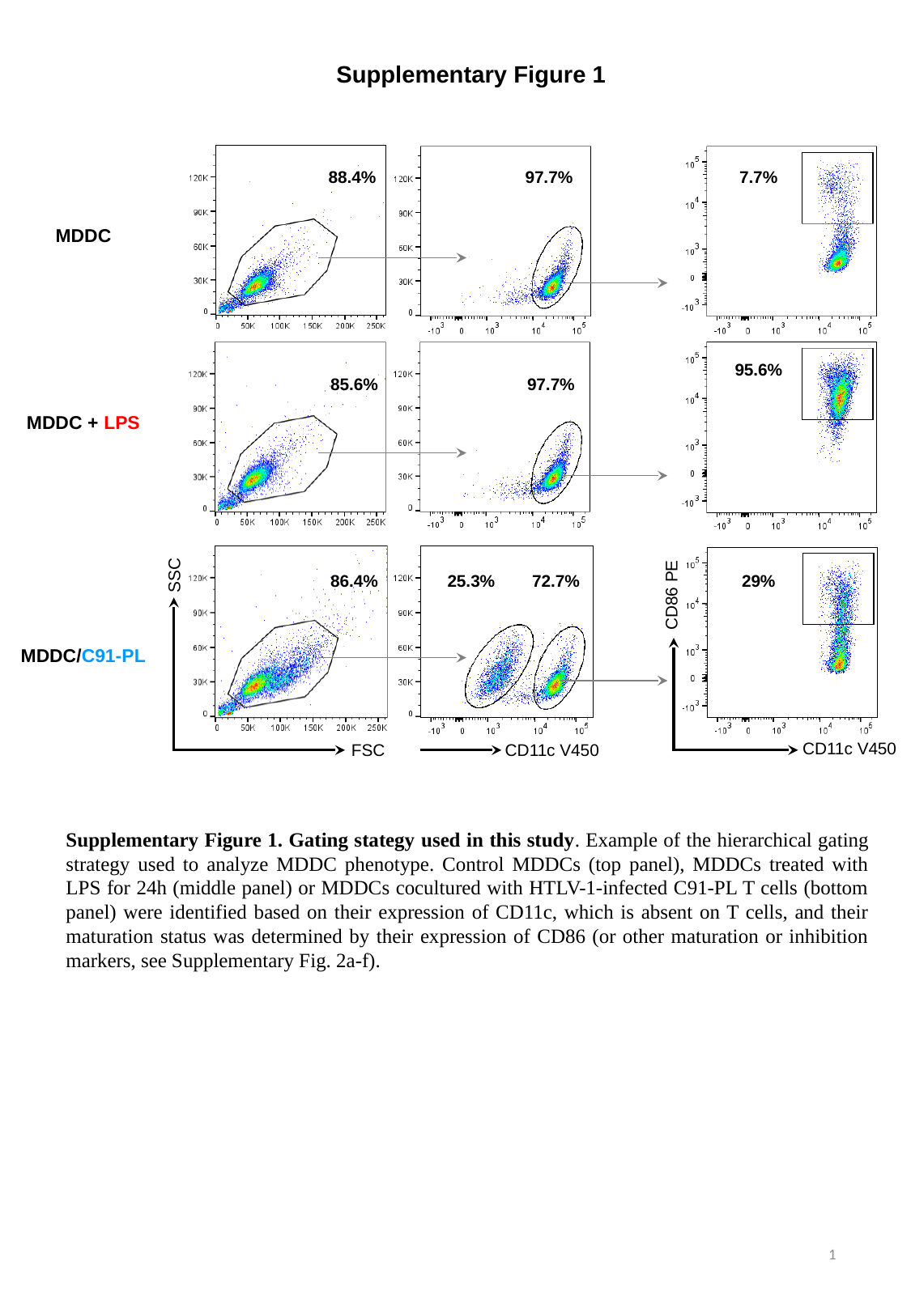

Supplementary Figure 1
88.4%
97.7%
7.7%
MDDC
95.6%
85.6%
97.7%
MDDC + LPS
SSC
29%
86.4%
25.3%
72.7%
CD86 PE
MDDC/C91-PL
CD11c V450
FSC
CD11c V450
Supplementary Figure 1. Gating stategy used in this study. Example of the hierarchical gating strategy used to analyze MDDC phenotype. Control MDDCs (top panel), MDDCs treated with LPS for 24h (middle panel) or MDDCs cocultured with HTLV-1-infected C91-PL T cells (bottom panel) were identified based on their expression of CD11c, which is absent on T cells, and their maturation status was determined by their expression of CD86 (or other maturation or inhibition markers, see Supplementary Fig. 2a-f).
1
