## Supplemental Figure 2 for "Peculiar transcriptional reprogramming with functional impairment of dendritic cells upon exposure to transformed HTLV-1-infected cells"

### Slide 1
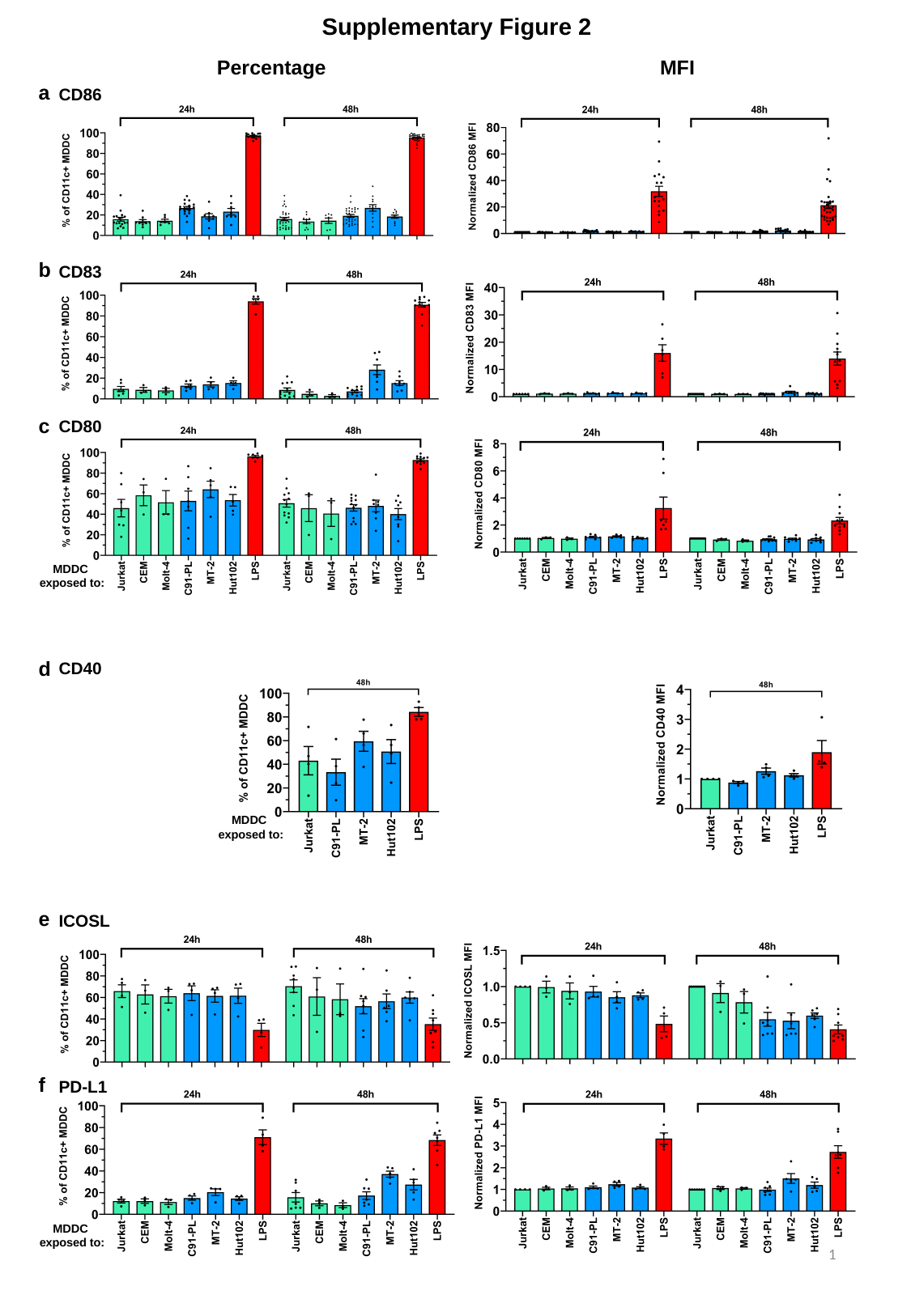

Supplementary Figure 2
MFI
Percentage
a
CD86
b
CD83
c
CD80
MDDC
exposed to:
d
CD40
MDDC
exposed to:
e
ICOSL
f
PD-L1
MDDC
exposed to:
1

### Slide 2
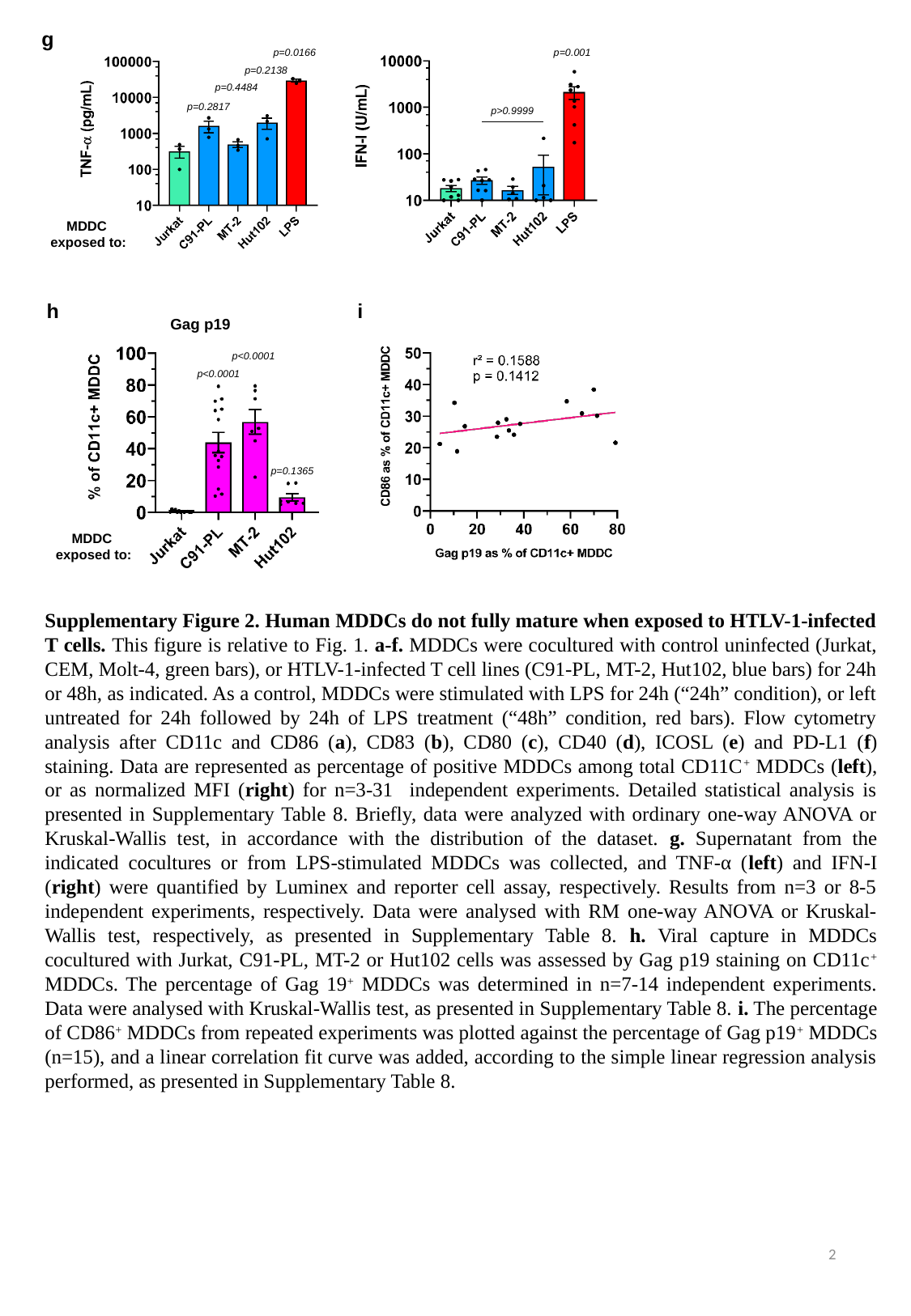

g
p=0.0166
p=0.001
p=0.2138
p=0.4484
p=0.2817
p>0.9999
MDDC
exposed to:
h
i
Gag p19
p<0.0001
p<0.0001
p=0.1365
MDDC
exposed to:
Supplementary Figure 2. Human MDDCs do not fully mature when exposed to HTLV-1-infected T cells. This figure is relative to Fig. 1. a-f. MDDCs were cocultured with control uninfected (Jurkat, CEM, Molt-4, green bars), or HTLV-1-infected T cell lines (C91-PL, MT-2, Hut102, blue bars) for 24h or 48h, as indicated. As a control, MDDCs were stimulated with LPS for 24h (“24h” condition), or left untreated for 24h followed by 24h of LPS treatment (“48h” condition, red bars). Flow cytometry analysis after CD11c and CD86 (a), CD83 (b), CD80 (c), CD40 (d), ICOSL (e) and PD-L1 (f) staining. Data are represented as percentage of positive MDDCs among total CD11C^+^ MDDCs (left), or as normalized MFI (right) for n=3-31 independent experiments. Detailed statistical analysis is presented in Supplementary Table 8. Briefly, data were analyzed with ordinary one-way ANOVA or Kruskal-Wallis test, in accordance with the distribution of the dataset. g. Supernatant from the indicated cocultures or from LPS-stimulated MDDCs was collected, and TNF-α (left) and IFN-I (right) were quantified by Luminex and reporter cell assay, respectively. Results from n=3 or 8-5 independent experiments, respectively. Data were analysed with RM one-way ANOVA or Kruskal-Wallis test, respectively, as presented in Supplementary Table 8. h. Viral capture in MDDCs cocultured with Jurkat, C91-PL, MT-2 or Hut102 cells was assessed by Gag p19 staining on CD11c^+^ MDDCs. The percentage of Gag 19^+^ MDDCs was determined in n=7-14 independent experiments. Data were analysed with Kruskal-Wallis test, as presented in Supplementary Table 8. i. The percentage of CD86^+^ MDDCs from repeated experiments was plotted against the percentage of Gag p19^+^ MDDCs (n=15), and a linear correlation fit curve was added, according to the simple linear regression analysis performed, as presented in Supplementary Table 8.
2
