## Supplemental Figure 3 for "Peculiar transcriptional reprogramming with functional impairment of dendritic cells upon exposure to transformed HTLV-1-infected cells"

### Slide 1
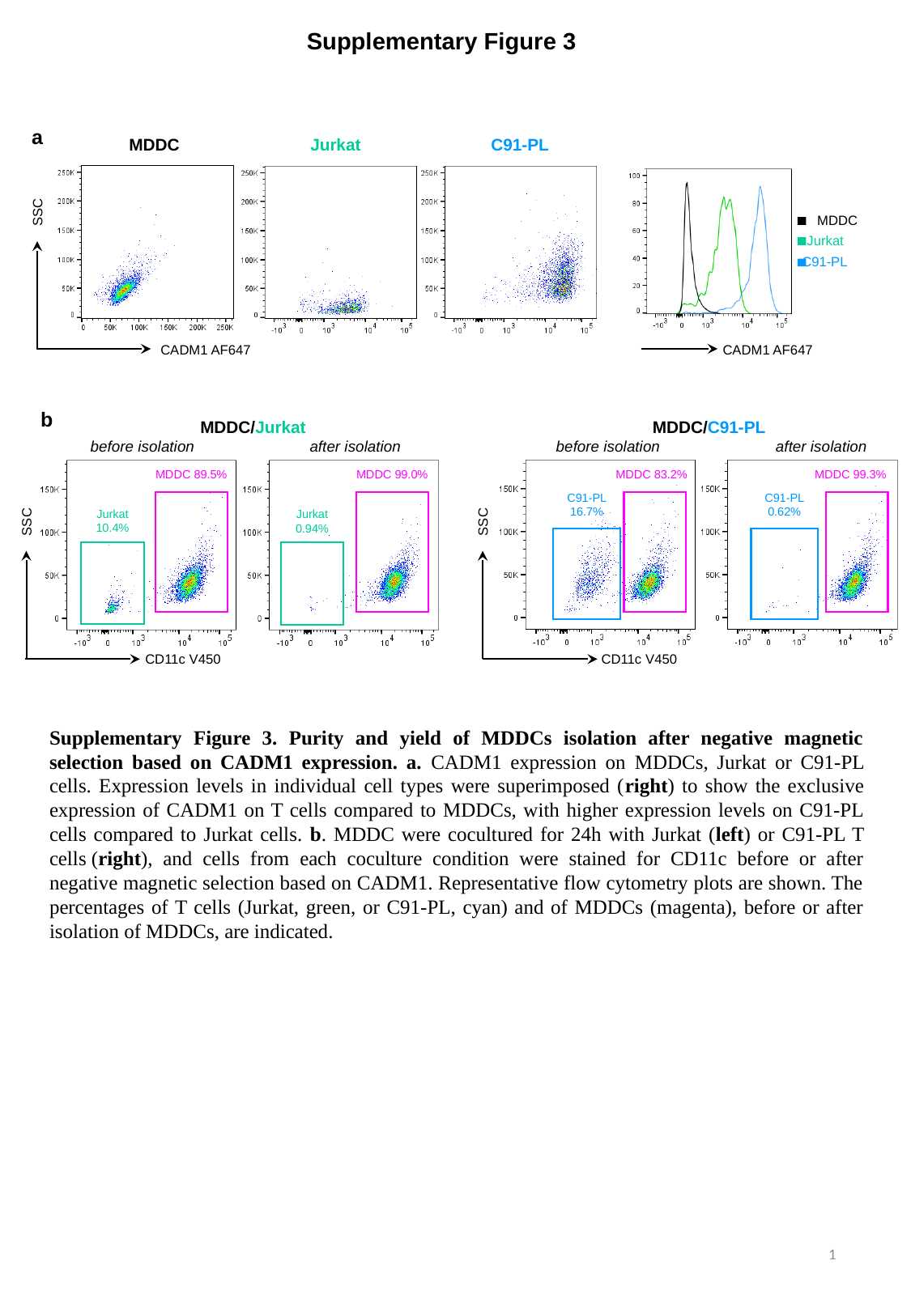

Supplementary Figure 3
a
MDDC
Jurkat
C91-PL
SSC
MDDC
C91-PL
Jurkat
CADM1 AF647
CADM1 AF647
b
MDDC/Jurkat
MDDC 89.5%
MDDC 99.0%
SSC
CD11c V450
Jurkat 10.4%
Jurkat 0.94%
MDDC/C91-PL
MDDC 83.2%
MDDC 99.3%
C91-PL 16.7%
C91-PL
0.62%
SSC
CD11c V450
before isolation
after isolation
before isolation
after isolation
Supplementary Figure 3. Purity and yield of MDDCs isolation after negative magnetic selection based on CADM1 expression. a. CADM1 expression on MDDCs, Jurkat or C91-PL cells. Expression levels in individual cell types were superimposed (right) to show the exclusive expression of CADM1 on T cells compared to MDDCs, with higher expression levels on C91-PL cells compared to Jurkat cells. b. MDDC were cocultured for 24h with Jurkat (left) or C91-PL T cells (right), and cells from each coculture condition were stained for CD11c before or after negative magnetic selection based on CADM1. Representative flow cytometry plots are shown. The percentages of T cells (Jurkat, green, or C91-PL, cyan) and of MDDCs (magenta), before or after isolation of MDDCs, are indicated.
1
