## Supplemental Figure 4 for "Peculiar transcriptional reprogramming with functional impairment of dendritic cells upon exposure to transformed HTLV-1-infected cells"

### Slide 1
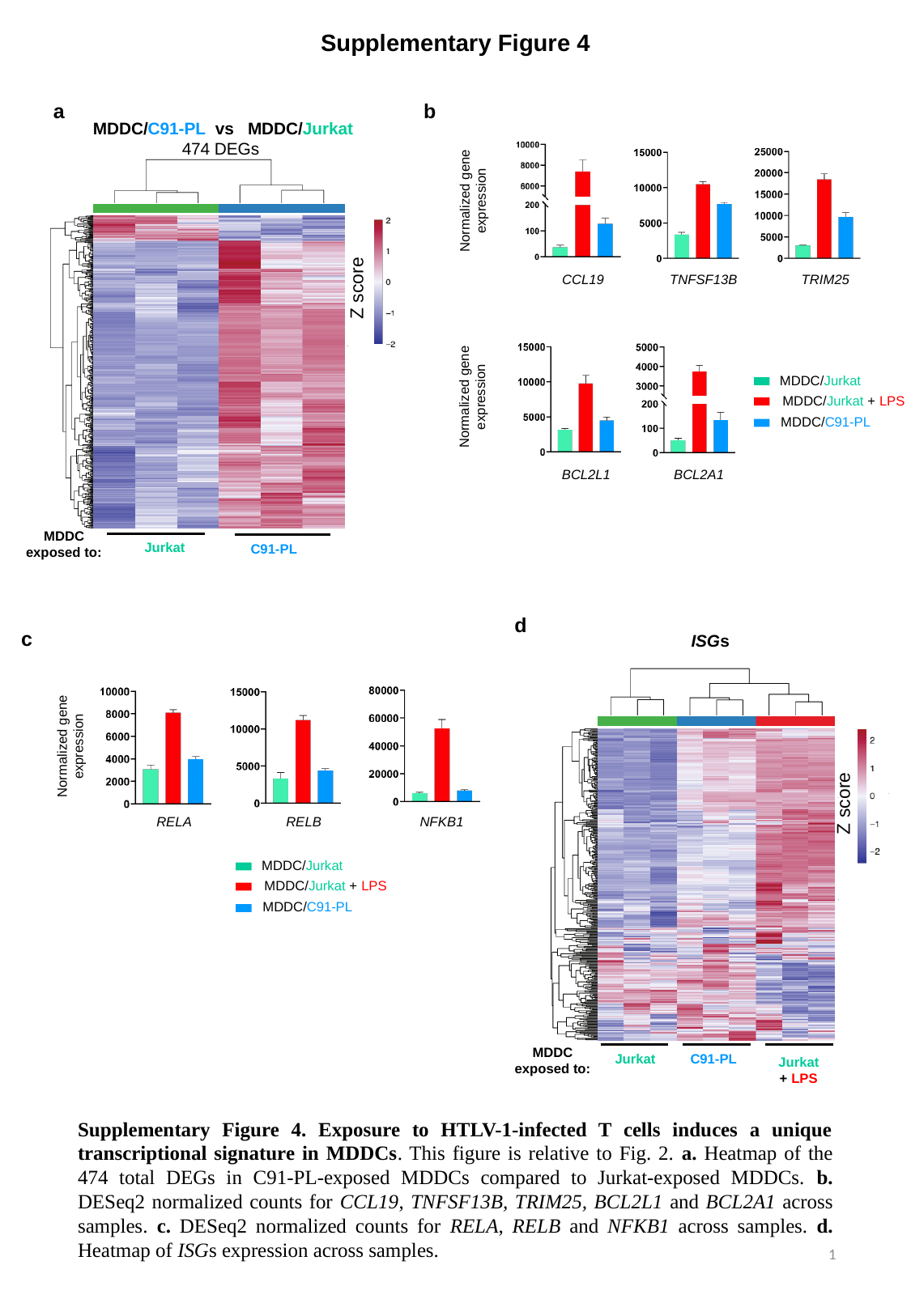

Supplementary Figure 4
a
b
MDDC/C91-PL vs MDDC/Jurkat
474 DEGs
Normalized gene expression
CCL19
TNFSF13B
TRIM25
Z score
MDDC/Jurkat
MDDC/C91-PL
MDDC/Jurkat + LPS
Normalized gene expression
BCL2L1
BCL2A1
MDDC
exposed to:
Jurkat
C91-PL
d
ISGs
Z score
MDDC
exposed to:
C91-PL
Jurkat
Jurkat
+ LPS
c
Normalized gene expression
RELA
RELB
NFKB1
MDDC/Jurkat
MDDC/C91-PL
MDDC/Jurkat + LPS
Supplementary Figure 4. Exposure to HTLV-1-infected T cells induces a unique transcriptional signature in MDDCs. This figure is relative to Fig. 2. a. Heatmap of the 474 total DEGs in C91-PL-exposed MDDCs compared to Jurkat-exposed MDDCs. b. DESeq2 normalized counts for CCL19, TNFSF13B, TRIM25, BCL2L1 and BCL2A1 across samples. c. DESeq2 normalized counts for RELA, RELB and NFKB1 across samples. d. Heatmap of ISGs expression across samples.
1
