## Supplemental Figure 5 for "Peculiar transcriptional reprogramming with functional impairment of dendritic cells upon exposure to transformed HTLV-1-infected cells"

### Slide 1
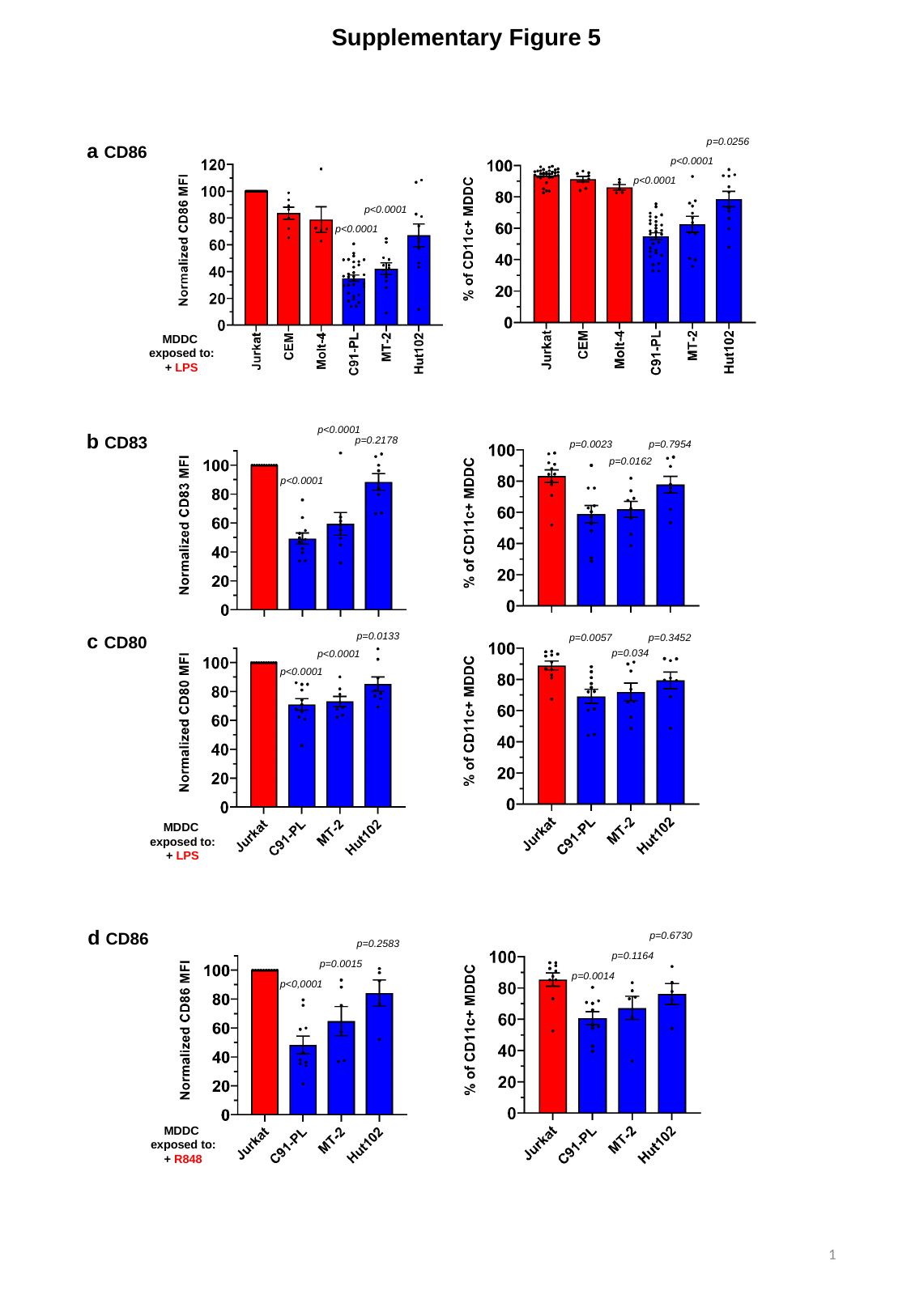

Supplementary Figure 5
a CD86
p=0.0256
p<0.0001
p<0.0001
p<0.0001
p<0.0001
MDDC
exposed to:
+ LPS
b CD83
p<0.0001
p=0.2178
p=0.0023
p=0.7954
p=0.0162
p<0.0001
c CD80
p=0.0133
p=0.0057
p=0.3452
p=0.034
p<0.0001
p<0.0001
MDDC
exposed to:
+ LPS
d CD86
p=0.6730
p=0.2583
p=0.1164
p=0.0015
p=0.0014
p<0,0001
MDDC
exposed to:
+ R848
1

### Slide 2
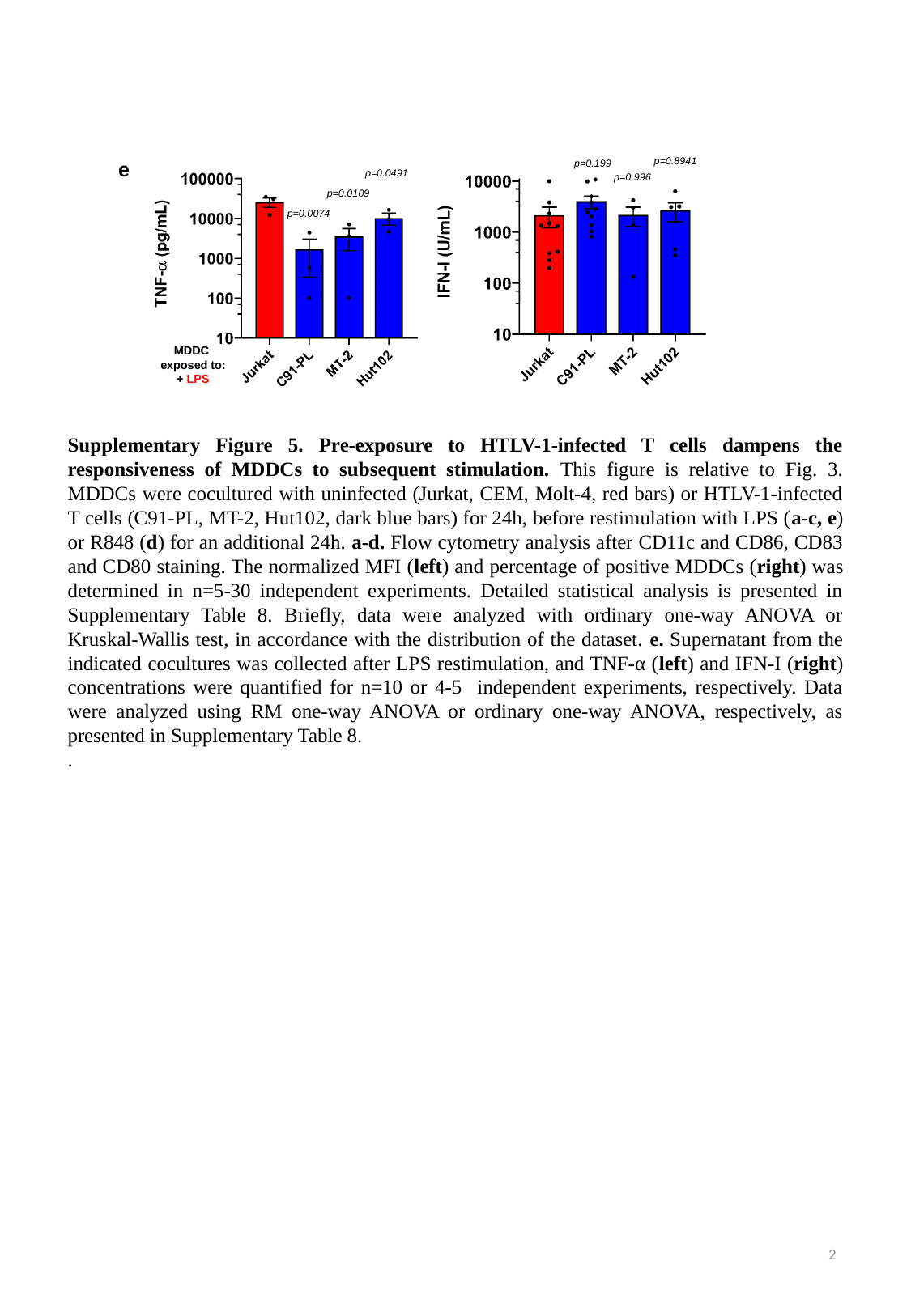

e
p=0.8941
p=0.199
p=0.0491
p=0.996
p=0.0109
p=0.0074
MDDC
exposed to:
+ LPS
Supplementary Figure 5. Pre-exposure to HTLV-1-infected T cells dampens the responsiveness of MDDCs to subsequent stimulation. This figure is relative to Fig. 3. MDDCs were cocultured with uninfected (Jurkat, CEM, Molt-4, red bars) or HTLV-1-infected T cells (C91-PL, MT-2, Hut102, dark blue bars) for 24h, before restimulation with LPS (a-c, e) or R848 (d) for an additional 24h. a-d. Flow cytometry analysis after CD11c and CD86, CD83 and CD80 staining. The normalized MFI (left) and percentage of positive MDDCs (right) was determined in n=5-30 independent experiments. Detailed statistical analysis is presented in Supplementary Table 8. Briefly, data were analyzed with ordinary one-way ANOVA or Kruskal-Wallis test, in accordance with the distribution of the dataset. e. Supernatant from the indicated cocultures was collected after LPS restimulation, and TNF-α (left) and IFN-I (right) concentrations were quantified for n=10 or 4-5 independent experiments, respectively. Data were analyzed using RM one-way ANOVA or ordinary one-way ANOVA, respectively, as presented in Supplementary Table 8.
.
2
