## Supplemental Figure 6 for "Peculiar transcriptional reprogramming with functional impairment of dendritic cells upon exposure to transformed HTLV-1-infected cells"

### Slide 1
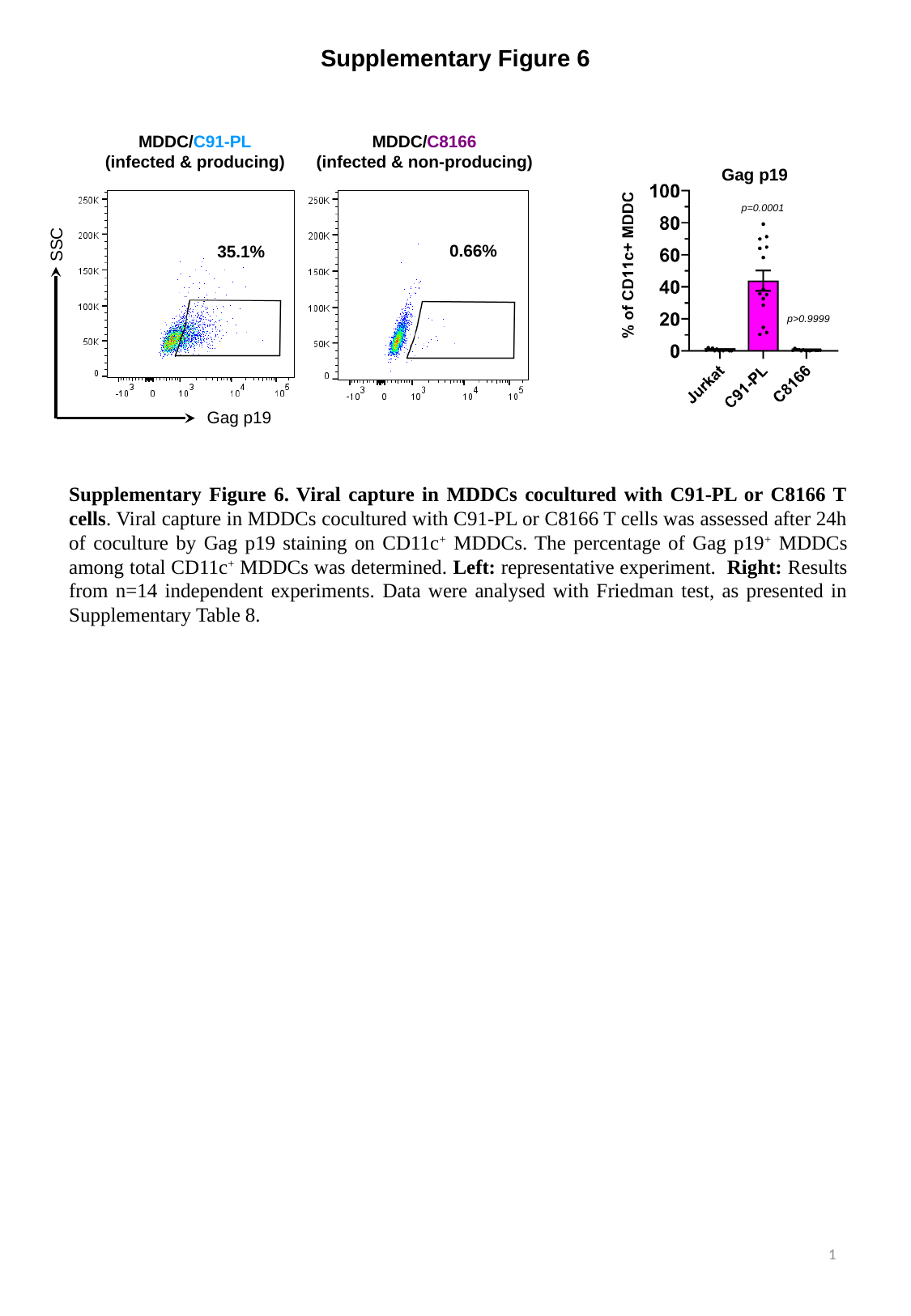

Supplementary Figure 6
MDDC/C91-PL
(infected & producing)
MDDC/C8166
(infected & non-producing)
SSC
0.66%
35.1%
Gag p19
Gag p19
p=0.0001
p>0.9999
Supplementary Figure 6. Viral capture in MDDCs cocultured with C91-PL or C8166 T cells. Viral capture in MDDCs cocultured with C91-PL or C8166 T cells was assessed after 24h of coculture by Gag p19 staining on CD11c^+^ MDDCs. The percentage of Gag p19^+^ MDDCs among total CD11c^+^ MDDCs was determined. Left: representative experiment. Right: Results from n=14 independent experiments. Data were analysed with Friedman test, as presented in Supplementary Table 8.
1
