## Supplemental Figure 7 for "Peculiar transcriptional reprogramming with functional impairment of dendritic cells upon exposure to transformed HTLV-1-infected cells"

### Slide 1
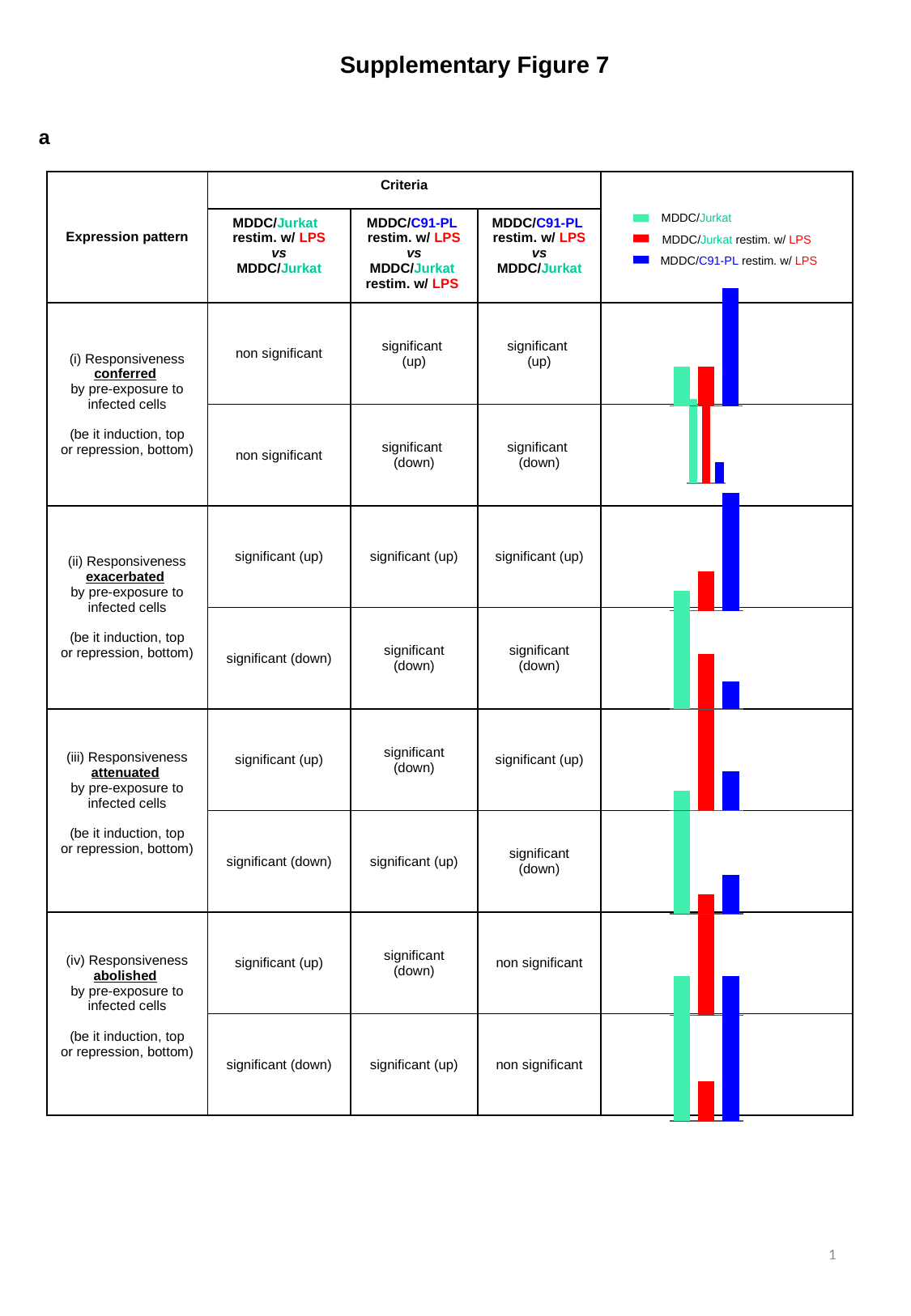

Supplementary Figure 7
a
| Expression pattern | Criteria | | |
| --- | --- | --- | --- |
| | MDDC/Jurkat restim. w/ LPS vs MDDC/Jurkat | MDDC/C91-PL restim. w/ LPS vs MDDC/Jurkat restim. w/ LPS | MDDC/C91-PL restim. w/ LPS vs MDDC/Jurkat |
| (i) Responsiveness conferred by pre-exposure to infected cells (be it induction, top or repression, bottom) | non significant | significant (up) | significant (up) |
| | non significant | significant (down) | significant (down) |
| (ii) Responsiveness exacerbated by pre-exposure to infected cells (be it induction, top or repression, bottom) | significant (up) | significant (up) | significant (up) |
| | significant (down) | significant (down) | significant (down) |
| (iii) Responsiveness attenuated by pre-exposure to infected cells (be it induction, top or repression, bottom) | significant (up) | significant (down) | significant (up) |
| | significant (down) | significant (up) | significant (down) |
| (iv) Responsiveness abolished by pre-exposure to infected cells (be it induction, top or repression, bottom) | significant (up) | significant (down) | non significant |
| | significant (down) | significant (up) | non significant |
MDDC/Jurkat
MDDC/Jurkat restim. w/ LPS
MDDC/C91-PL restim. w/ LPS
#### Chart
| Category | |
|---|---|
| DC/JK | 1.0 |
| DC/JK LPS | 1.0 |
| DC/C91/LPS | 3.0 |
#### Chart
| Category | |
|---|---|
| DC/JK | 2.0 |
| DC/JK LPS | 2.0 |
| DC/C91/LPS | 0.5 |
#### Chart
| Category | |
|---|---|
| DC/JK | 0.5 |
| DC/JK LPS | 1.0 |
| DC/C91/LPS | 3.0 |
#### Chart
| Category | |
|---|---|
| DC/JK | 2.0 |
| DC/JK LPS | 1.0 |
| DC/C91/LPS | 0.5 |
#### Chart
| Category | |
|---|---|
| DC/JK | 0.5 |
| DC/JK LPS | 3.0 |
| DC/C91/LPS | 1.0 |
#### Chart
| Category | |
|---|---|
| DC/JK | 3.0 |
| DC/JK LPS | 0.5 |
| DC/C91/LPS | 1.0 |
#### Chart
| Category | |
|---|---|
| DC/JK | 1.0 |
| DC/JK LPS | 3.0 |
| DC/C91/LPS | 1.0 |
#### Chart
| Category | |
|---|---|
| DC/JK | 3.0 |
| DC/JK LPS | 1.0 |
| DC/C91/LPS | 3.0 |1

### Slide 2
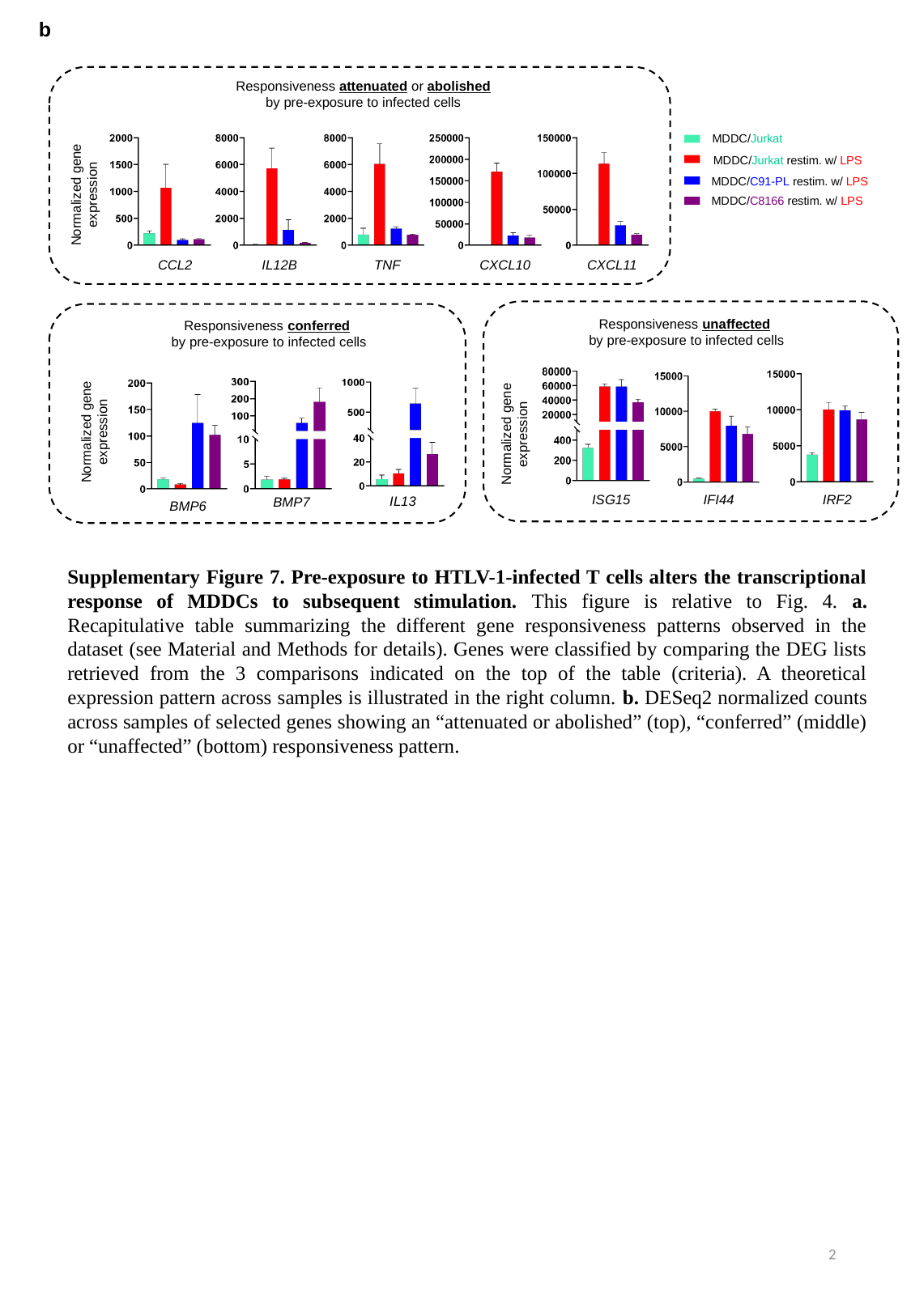

b
Responsiveness attenuated or abolished by pre-exposure to infected cells
Normalized gene expression
CCL2
CXCL10
CXCL11
IL12B
TNF
MDDC/Jurkat
MDDC/Jurkat restim. w/ LPS
MDDC/C91-PL restim. w/ LPS
MDDC/C8166 restim. w/ LPS
Responsiveness conferred
by pre-exposure to infected cells
IL13
BMP6
BMP7
Normalized gene expression
Responsiveness unaffected
by pre-exposure to infected cells
Normalized gene expression
ISG15
IFI44
IRF2
Supplementary Figure 7. Pre-exposure to HTLV-1-infected T cells alters the transcriptional response of MDDCs to subsequent stimulation. This figure is relative to Fig. 4. a. Recapitulative table summarizing the different gene responsiveness patterns observed in the dataset (see Material and Methods for details). Genes were classified by comparing the DEG lists retrieved from the 3 comparisons indicated on the top of the table (criteria). A theoretical expression pattern across samples is illustrated in the right column. b. DESeq2 normalized counts across samples of selected genes showing an “attenuated or abolished” (top), “conferred” (middle) or “unaffected” (bottom) responsiveness pattern.
2
