## Supplemental Figure 8 for "Peculiar transcriptional reprogramming with functional impairment of dendritic cells upon exposure to transformed HTLV-1-infected cells"

### Slide 1
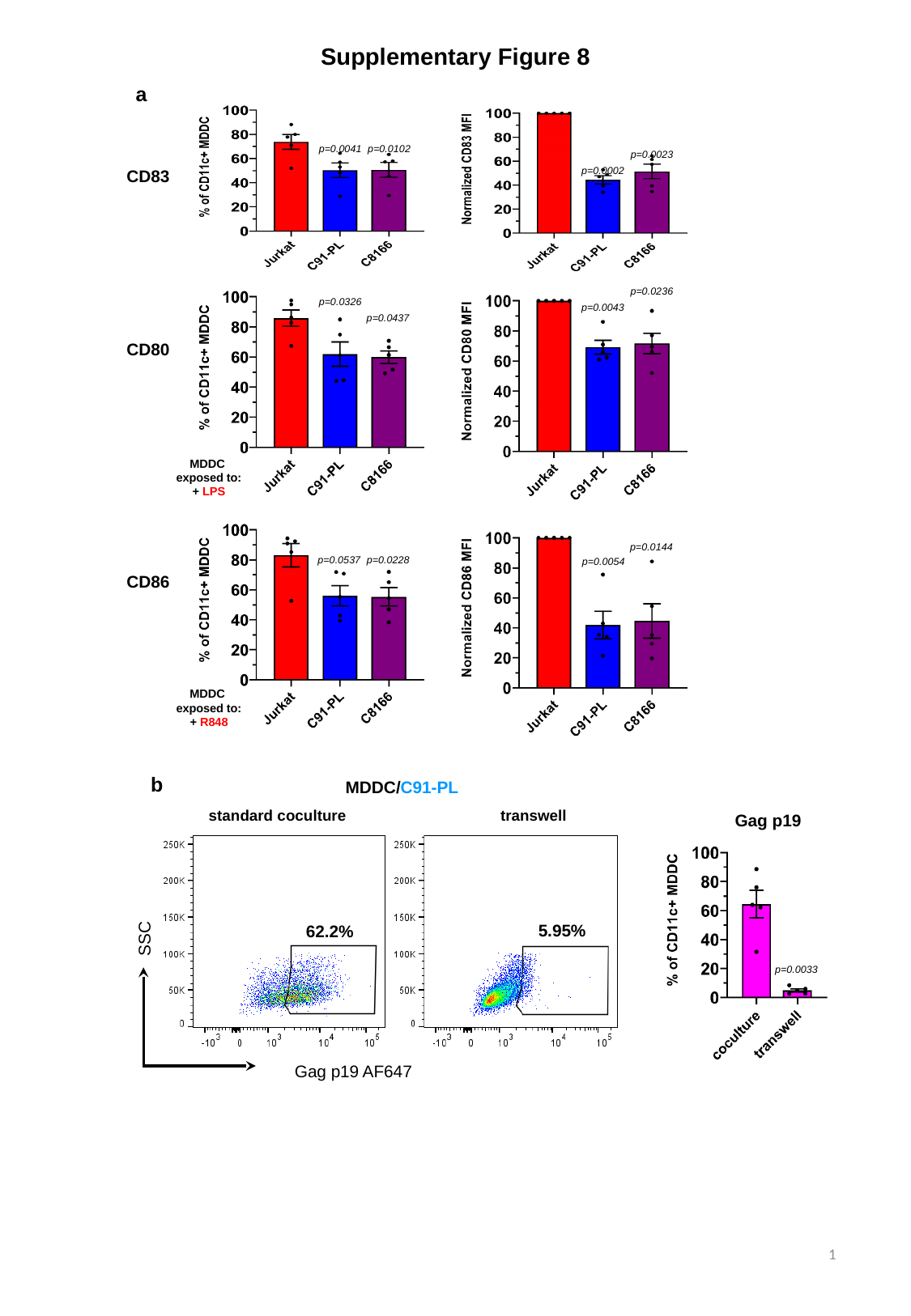

Supplementary Figure 8
a
CD83
CD80
MDDC
exposed to:
+ LPS
CD86
MDDC
exposed to:
+ R848
p=0.0041
p=0.0102
p=0.0023
p=0.0002
p=0.0236
p=0.0326
p=0.0043
p=0.0437
p=0.0144
p=0.0537
p=0.0228
p=0.0054
b
MDDC/C91-PL
standard coculture
transwell
SSC
5.95%
62.2%
Gag p19 AF647
Gag p19
p=0.0033
1

### Slide 2
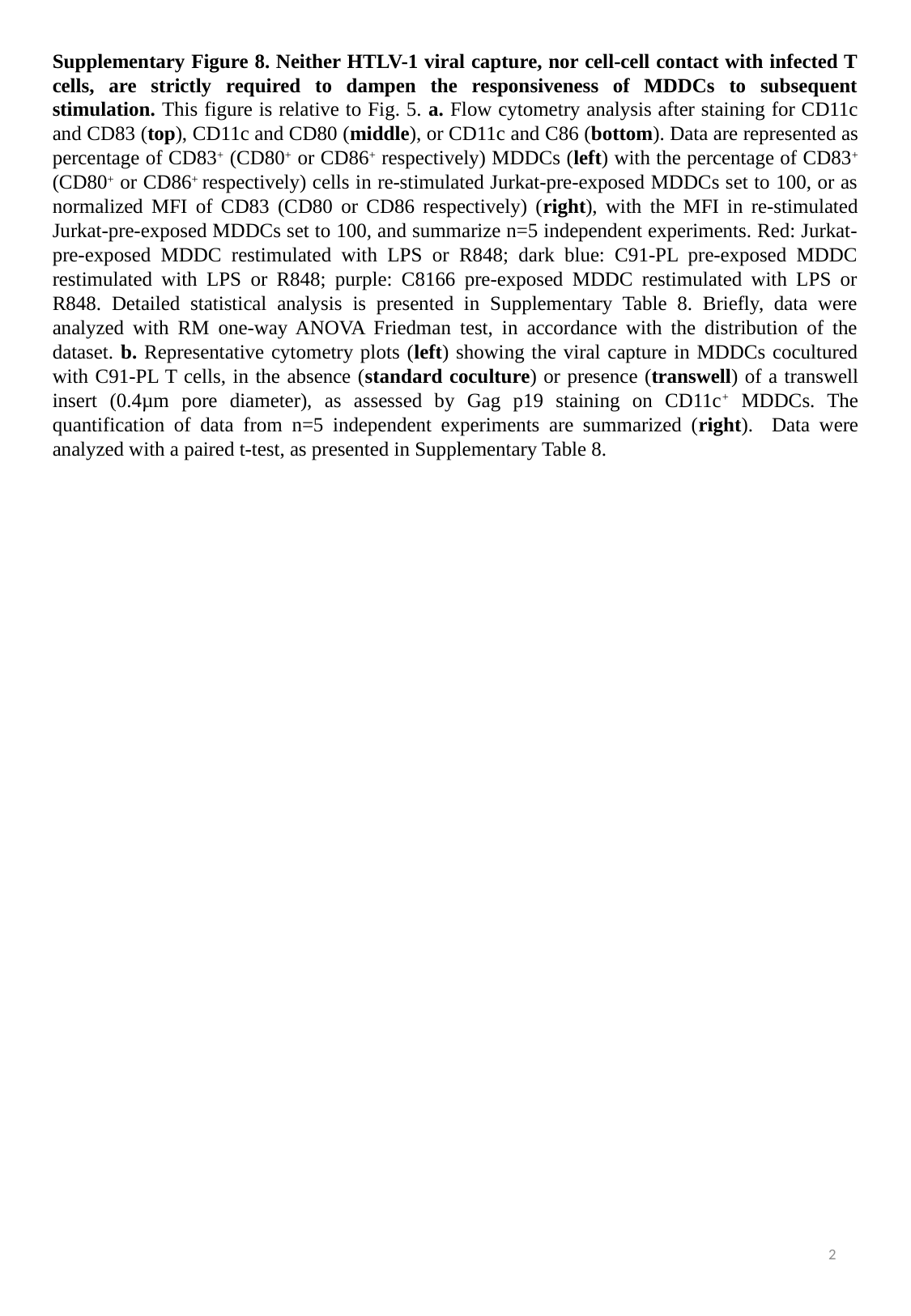

Supplementary Figure 8. Neither HTLV-1 viral capture, nor cell-cell contact with infected T cells, are strictly required to dampen the responsiveness of MDDCs to subsequent stimulation. This figure is relative to Fig. 5. a. Flow cytometry analysis after staining for CD11c and CD83 (top), CD11c and CD80 (middle), or CD11c and C86 (bottom). Data are represented as percentage of CD83^+^ (CD80^+^ or CD86^+^ respectively) MDDCs (left) with the percentage of CD83^+^ (CD80^+^ or CD86^+^ respectively) cells in re-stimulated Jurkat-pre-exposed MDDCs set to 100, or as normalized MFI of CD83 (CD80 or CD86 respectively) (right), with the MFI in re-stimulated Jurkat-pre-exposed MDDCs set to 100, and summarize n=5 independent experiments. Red: Jurkat-pre-exposed MDDC restimulated with LPS or R848; dark blue: C91-PL pre-exposed MDDC restimulated with LPS or R848; purple: C8166 pre-exposed MDDC restimulated with LPS or R848. Detailed statistical analysis is presented in Supplementary Table 8. Briefly, data were analyzed with RM one-way ANOVA Friedman test, in accordance with the distribution of the dataset. b. Representative cytometry plots (left) showing the viral capture in MDDCs cocultured with C91-PL T cells, in the absence (standard coculture) or presence (transwell) of a transwell insert (0.4µm pore diameter), as assessed by Gag p19 staining on CD11c^+^ MDDCs. The quantification of data from n=5 independent experiments are summarized (right). Data were analyzed with a paired t-test, as presented in Supplementary Table 8.
2
