## Supplemental Figure 9 for "Peculiar transcriptional reprogramming with functional impairment of dendritic cells upon exposure to transformed HTLV-1-infected cells"

### Slide 1
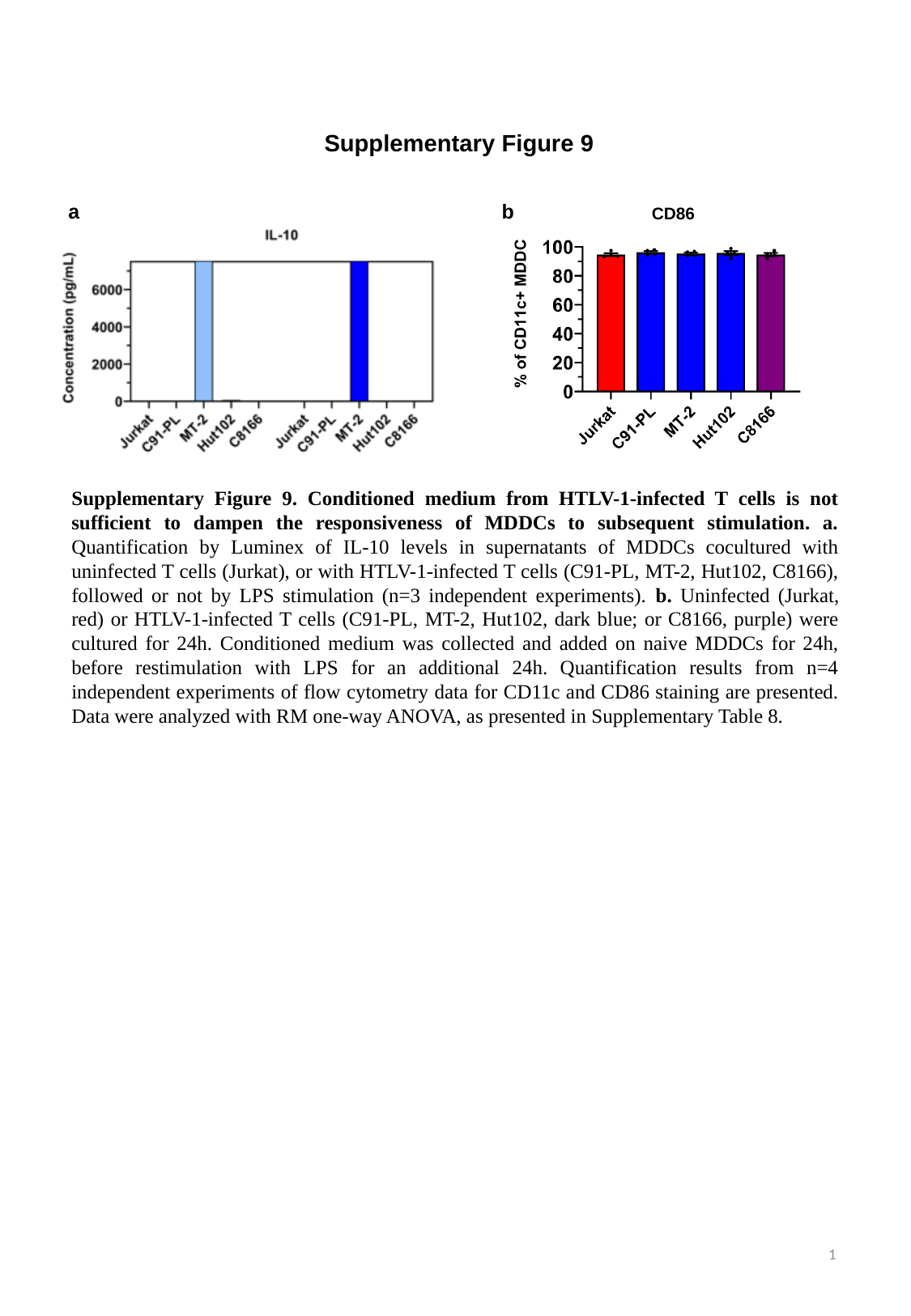

Supplementary Figure 9
a
b
CD86
Supplementary Figure 9. Conditioned medium from HTLV-1-infected T cells is not sufficient to dampen the responsiveness of MDDCs to subsequent stimulation. a. Quantification by Luminex of IL-10 levels in supernatants of MDDCs cocultured with uninfected T cells (Jurkat), or with HTLV-1-infected T cells (C91-PL, MT-2, Hut102, C8166), followed or not by LPS stimulation (n=3 independent experiments). b. Uninfected (Jurkat, red) or HTLV-1-infected T cells (C91-PL, MT-2, Hut102, dark blue; or C8166, purple) were cultured for 24h. Conditioned medium was collected and added on naive MDDCs for 24h, before restimulation with LPS for an additional 24h. Quantification results from n=4 independent experiments of flow cytometry data for CD11c and CD86 staining are presented. Data were analyzed with RM one-way ANOVA, as presented in Supplementary Table 8.
1
